# “Regulation of Obesity and Fatty Liver by Moringa oleifera: Insights into Inflammatory Pathways”

**DOI:** 10.1101/2024.04.28.591562

**Authors:** Nihal A. Ibrahim, Manal A. Buabeid, El-Shaimaa A. Arafa, Ghulam Murtaza

## Abstract

Obesity and fatty liver are relatively benign states but continued inflammatory stress and its metabolic implications turn them into one of the most devastating diseases of humankind. Generally, obesity and fatty liver precede diabetes mellitus, cardiovascular problems and malignant growths. The present research aimed to explore the efficacy of methanolic extract of *Moringa Olifera* (*Me*.*MO*) for the management of obesity and fatty liver and related inflammatory state that prime the body for devastating effects. A series of *in-vitro* and *in-vivo* studies were employed. Data from HPLC analysis confirmed the presence of flavonoids and phenolic acids. Rats were fed on either normal diet (ND) or high fat diet (HFD and streptozocin (STZ) in the presence or absence of *Me*.*Mo* (250 mg/kg & 500 mg/kg) or metformin (70 mg/kg). Findings showed that rats received 500 mg/kg *Me*.*MO* showed a significant (*p* > 0.01) decrease in body weights, liver weights, and plasma glucose level. Laboratory data exhibited a significant (*p* < 0.05) inhibitory effect on *Me*.*MO* on pro-inflammatory mediators (IL-1B and TNF) and caused a sharp increase in anti-inflammatory cytokines levels (IL-10, IL-6 and COX-2) in all treatment groups. Histopathological analysis exhibited no structural and functional alteration in the liver and adipose tissues. Altogether, *Me*.*MO* ameliorates experimentally induced obesity accompanying fatty liver and inflammatory stress. However, further investigations are still needed to confirm the safety and efficacy of *Moringaβ olifera* (MO) for clinical application.

## 1. Introduction

The view that obesity and related metabolic complications are not merely lipid overload conditions but a chronic low-grade inflammatory state with metabolic repercussions has largely changed our understanding of the disease. It is now understood that most obese states incite the immune system [1] that themselves undergo metabolic changes pressed by fats [2-4] to initiate a vicious immunometabolic dysregulation cycle [5, 6], often culminating in complicated disorders [7, 8]. Obesity with a low-grade inflammatory state is often accompanied by fatty liver [9], serum lipid dysregulations [10] and fatal cardiovascular complications [11]. Currently, there are five FDA-approved drugs for chronic obesity namely orlistat, phentermine-topiramate, naltrexone-bupropion, liraglutide, and semaglutide [12] but these medications primarily focus on suppressing appetite by targeting GLP-1 [13] or work by reducing the absorption of fat from intestine [14]. The lack of anti-inflammatory actions, inability to systemically regulate lipids, varied pharmacokinetic profile and spectrum of adverse drug reactions leave a room for better future choices.

*Moringa Olifera* (MO) is a tree that belongs to the family *Moringaceae* and is commonly known as the “Miracle Tree”. It is native to Asia and Africa and widely grown in tropical and subtropical climates. The leaves, roots, pods, flowers and seeds of MO possess both nutritional as well as medicinal properties [15]. Traditionally, MO has been employed extensively for the treatment of malaria, typhoid, hematological, cardiovascular and gastrointestinal disorders [16]. Several studies have demonstrated the presence of secondary metabolites vitamins (A, B, C), alkaloids, glycosides, flavonoids, tannins, saponins and terpenoids in *MO* tree [17]. Some flavonoids (myricetin, quercetin, kaempferol, isorhamnetin and rutin) present in MO leaves and seeds showed excellent antimicrobial and antiproliferative (inhibits S and G2M pathways) and anti-apoptotic effects (downregulate nuclear factor kappa-B) [18, 19]. Recently, various pre-clinical and clinical studies have evidenced multiple biological activities of MO including gastro-protectant, hypotensive, antidiabetic, hepatoprotective, antimicrobial, antihyperlipidemic, anticancer and anti-inflammatory. It has also been exhibited therapeutic effects on renal injury, and thyroid hormone regulation [20]. The anti-inflammatory potential of MO are already reported in cancer [21] and renal damage [22] but how it exerts its anti-inflammatory actions under lipid overload states is yet to be discovered.

This study aims to investigate the MO potential in obesity-induced fatty liver and accompanied inflammatory state. This study forms the basis for further experimentation of MO on the immunometabolic axis, which is pivotal for holistic obesity treatment.

## 2. Materials and Methods

### 2.1. Collection and Extraction

The whole plant was collected from Benessere Health Company (BHC), Multan, Pakistan in June 2020 and authenticated by Dr. Altaf Dashti and voucher specimen 3456AS was deposited at the herbarium of the Department of Botany, University of Punjab, Lahore, Pakistan. Firstly, the plant was dried under shade for 14 days then crushed into fine powder and soaked in analytical grade methanol for maceration. The rotary evaporator was under for excessive solvent evaporation under reduced pressure and temperature (45°C). The percentage yield of extract was calculated to be 5% w/w and labeled as Me*-MO*.

### 2.2. *In-vivo* activities

#### 2.2.1 Animals and treatment

Twenty-five Wistar albino rats (10-12 weeks old) were purchased from the animal house of Riphah International University Lahore. Animals were kept in polypropylene cages under 12 h light/dark cycle with the provision of sufficient food and water supply. All study protocols had been followed as authorized by the Animals Ethical Committee of COMSATS University Islamabad, Lahore campus. Animals were acclimatized for one week. Animals were divided into five corresponding groups (n=5). Group-I served as the control group while group-II was administered high fat diet (HFD). Remaining groups either received MO at 250 mg/kg and 500 mg/kg or received metformin (70 mg/kg in 5% CMC). All of the groups were fed on HFD (24g% fat + 24g% protein + 41g% carbohydrates) for 12 weeks except group-I that received normal diet (ND).

#### 2.2.2. Induction of obesity and lipid dysregulation

At the start of the 1^st^ week, the body weight and blood glucose level of all animals were measured with the help of glucose oxidase reagent strips (Accu-Chek^®^). The diabetogenic agent streptozotocin (STZ) (30 mg/kg) was injected intraperitoneally in addition to HFD to all animals except group-I [23]. After 48 hours, glucose level was again measured and defined doses of plant extract as well as standard was administrated for the period of the next 8 weeks.

#### 2.2.3. Physical parameters

The glucose level and body weight of all animals were measured after a constant interval (1 week). All rats were maintained on HFD throughout the study except the normal group. On the final day, animals were slaughtered, and blood was drawn for biochemical and hematological analysis. Changes in weight of liver and adipose tissue were also estimated through Microsoft excel software®.

#### 2.2.4. Biochemical parameters

Serum analysis was performed to analyze the effect of *Me*.*MO* on random blood glucose, low-density lipoproteins (LDL), high-density lipoproteins (HDL), triglycerides (TG), cholesterol, alanine aminotransferase (ALT) aspartate aminotransferase (AST), and triglycerides (TGs) by using commercial kits as previously described [24].

#### 2.2.5. Inflammatory parameters

The concentration of various inflammatory biomarkers including TNF-α, IL-6, IL-1β, NF-kβ and IL-10 in serum was determined through ELISA kits [25].

#### 2.2.6. Histopathological analysis

Liver and adipose tissues were excised and preserved in formalin solution for histopathological examination. Slides were prepared by using hematoxylin and eosin dyes and analyzed under a microscope for any pathological change [26].

### 2.3. *In-vitro* analysis

#### 2.3.1. High-Performance Liquid Chromatography (HPLC)

To quantify phytoconstituents in *Me*.*MO*, high-pressure liquid chromatography was performed. By using two different types of mobile phases having a composition as phase-A (water and acetic acid in a ratio of 94:6 and pH = 2.27) and phase-B including acetonitrile 15% from 0-15 minutes followed by 45% from 15-30 minutes and 100% from 30-45 minutes. Shim-pack HPLC (CLC-ODS; C-18; Shimadzu, Japan) column having length/height dimensions of 25 cm × 4.6 mm and 5 µm diameter was used to isolate the compounds. Analysis of samples was done with help of an ultraviolet detector (280 nm wavelength). In the end, a chromatogram was drawn between voltage (x-axis) and time (y-axis) units respectively, and compared with standards [27].

### 2.4 Statistical analysis

The analysis of data was done through Graph-Pad Prism Software and Microsoft Excel. Results are presented as mean ± SEM after applying the one-way analysis of variance. The value of *“p”* below 0.05 is assumed to be significant.

## 3. Results

### 3.1 HPLC analysis

Data in Table 1 shows five compounds including quercetin, vanillic acid, chlorogenic acid, synergic acid and m-coumaric acid obtained after HPLC analysis of *Me*.*MO*. The retention time, percentage areas and concentration of all phenolic acids in parts per million are also presented. Moreover, peaks of all identified compounds are shown in the chromatogram (Figure 1).

**Table 1:**
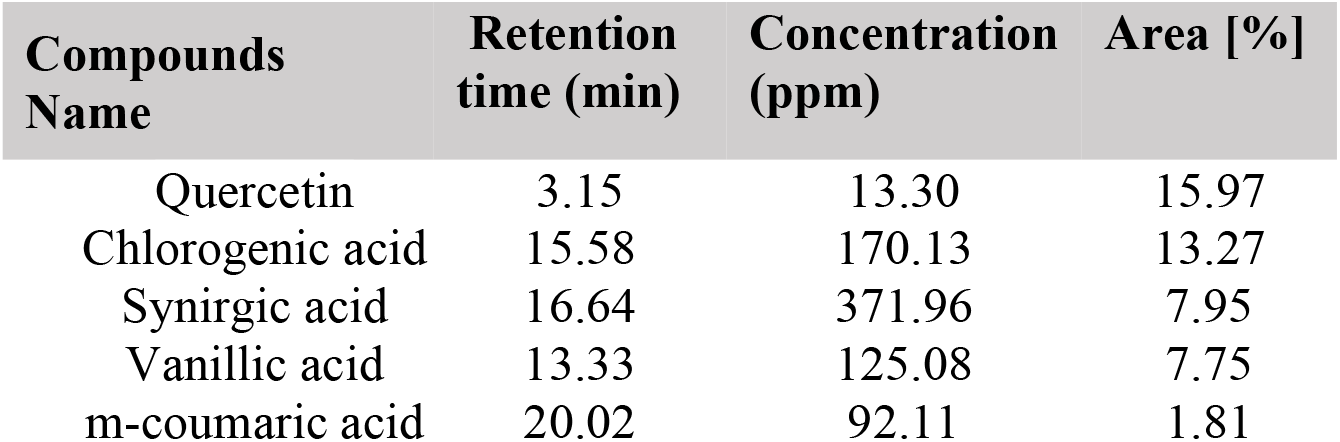
Compounds identified in HPLC.

**Figure 1:**
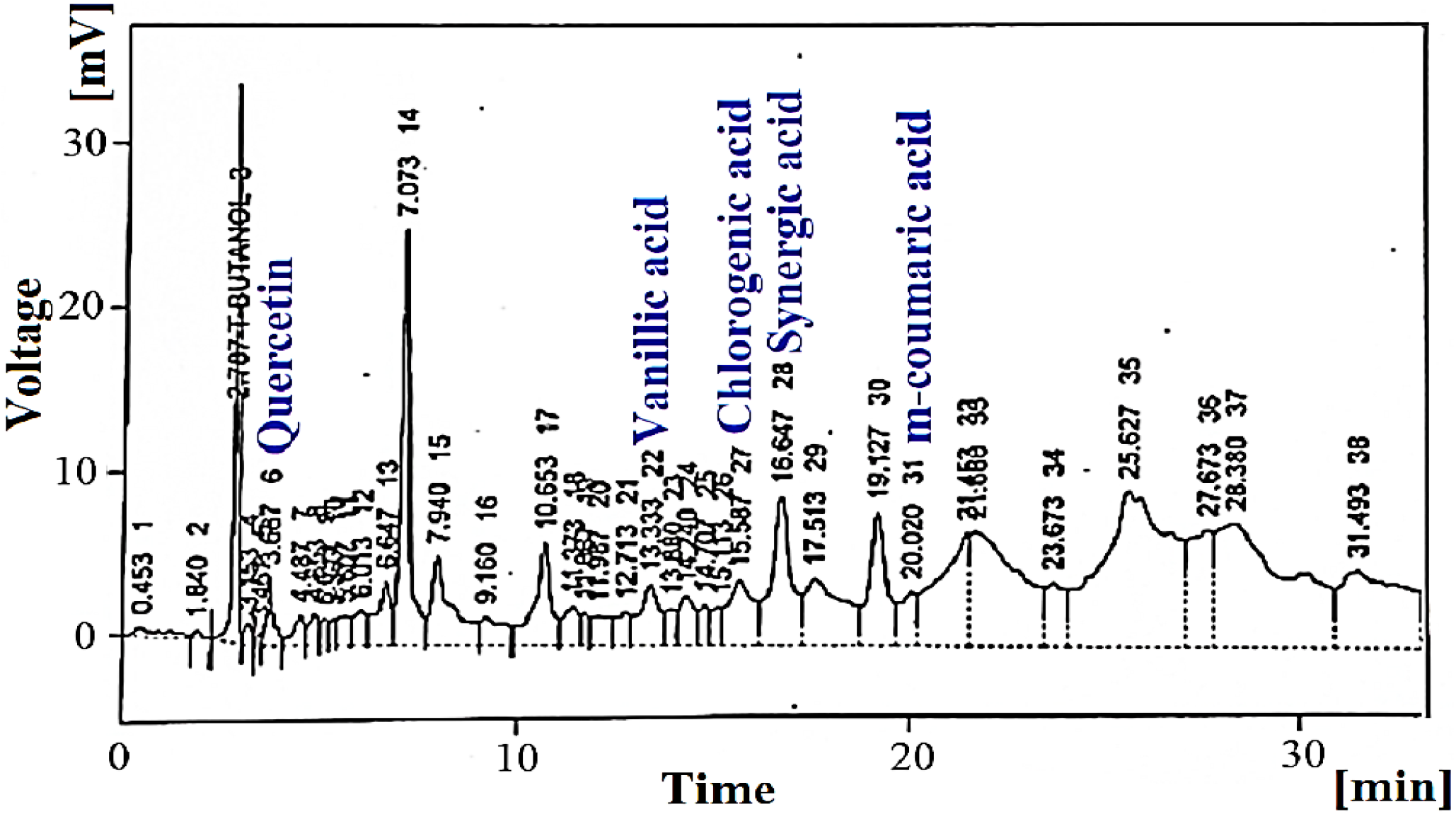
HPLC chromatogram of *Me*.*MO*

### 3.2 *Me.Mo* reduces body weights and liver weights

Figure 2 and figure 3 represent the effect of *Me*.*MO* on body weight of animals before, during and after the study. On zero-week, no significant (*p* > 0.05) difference was observed in body weights of all groups statistically. While at 4^th^ week, the weight of animals drastically rose after the administration of HFD. Statistical analysis showed a significant (*p* < 0.001) increase in the weight of animals after receiving the HFD. On the 12^th^ week, weight variation for control, model (HFD), *Me*.*MO* 250 mg/kg, *Me*.*MO* 500 mg/kg and metformin (70 mg/kg) was calculated to be 205.0 ± 0.55, 291.0 ± 1.00, 206.0 ± 0.15, 215.0 ± 0.30 and 186.0 ± 0.90 respectively. In addition, macroscopic analysis with a naked eye also showed a marked increase and decrease in body weight of rats at the initial (4^th^ week) and final stage (12^th^ week) of the study (Figure 3). Similarly, liver weight was also increased in HFD induced group while administration of *Me*.*Mo* significantly reversed the raised liver weights (Figure 4).

**Figure 2:**
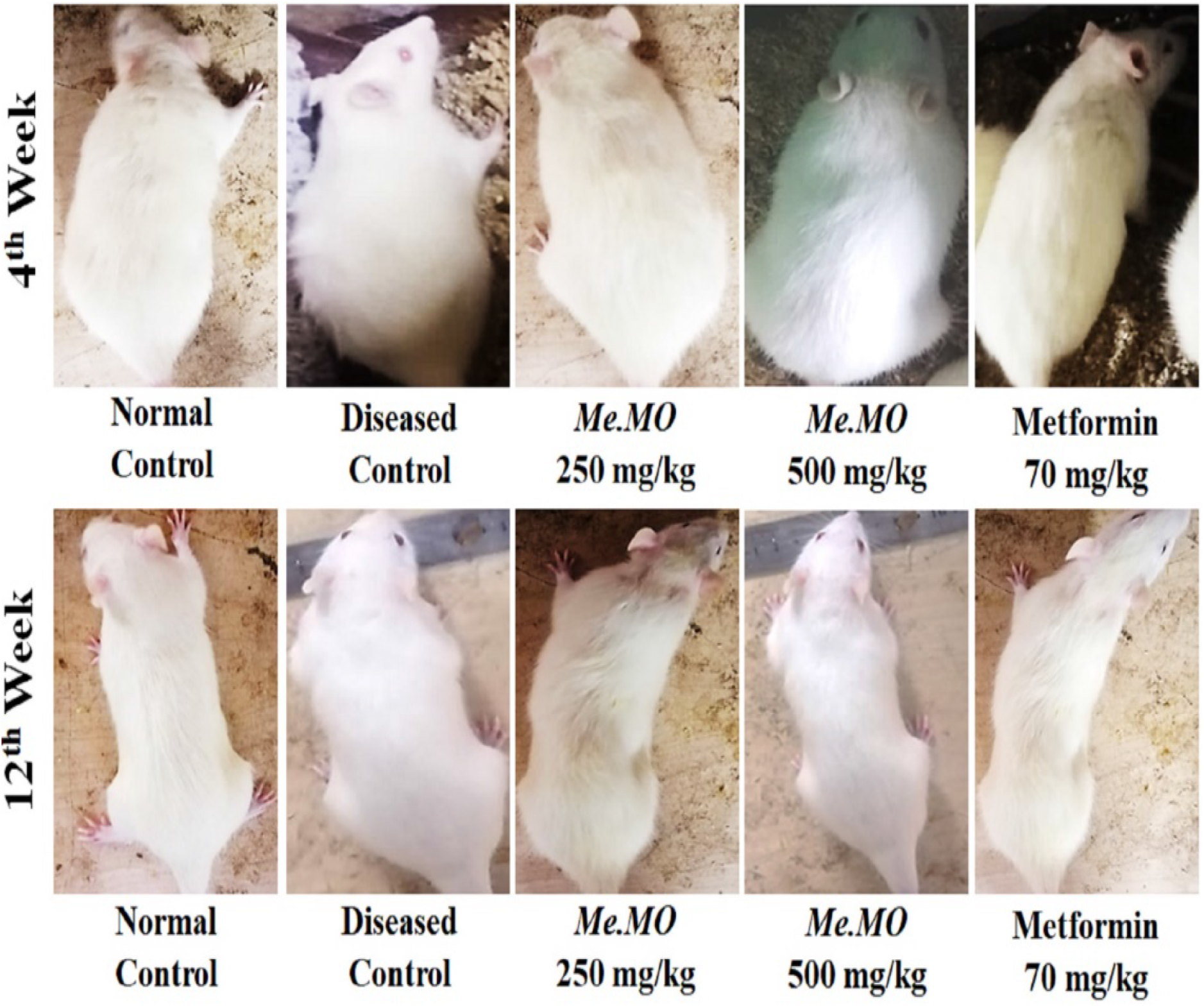
Macroscopic analysis of body weight variations on 0, 4^th^ and 12^th^ week in different study groups shown in representative pictographic form. A visitor, not knowing the background of study, chose subjects for macroscopic analysis randomly. Animals were weigh after the pictures were taken and found consistent with the macroscopic analysis.

**Figure 3:**
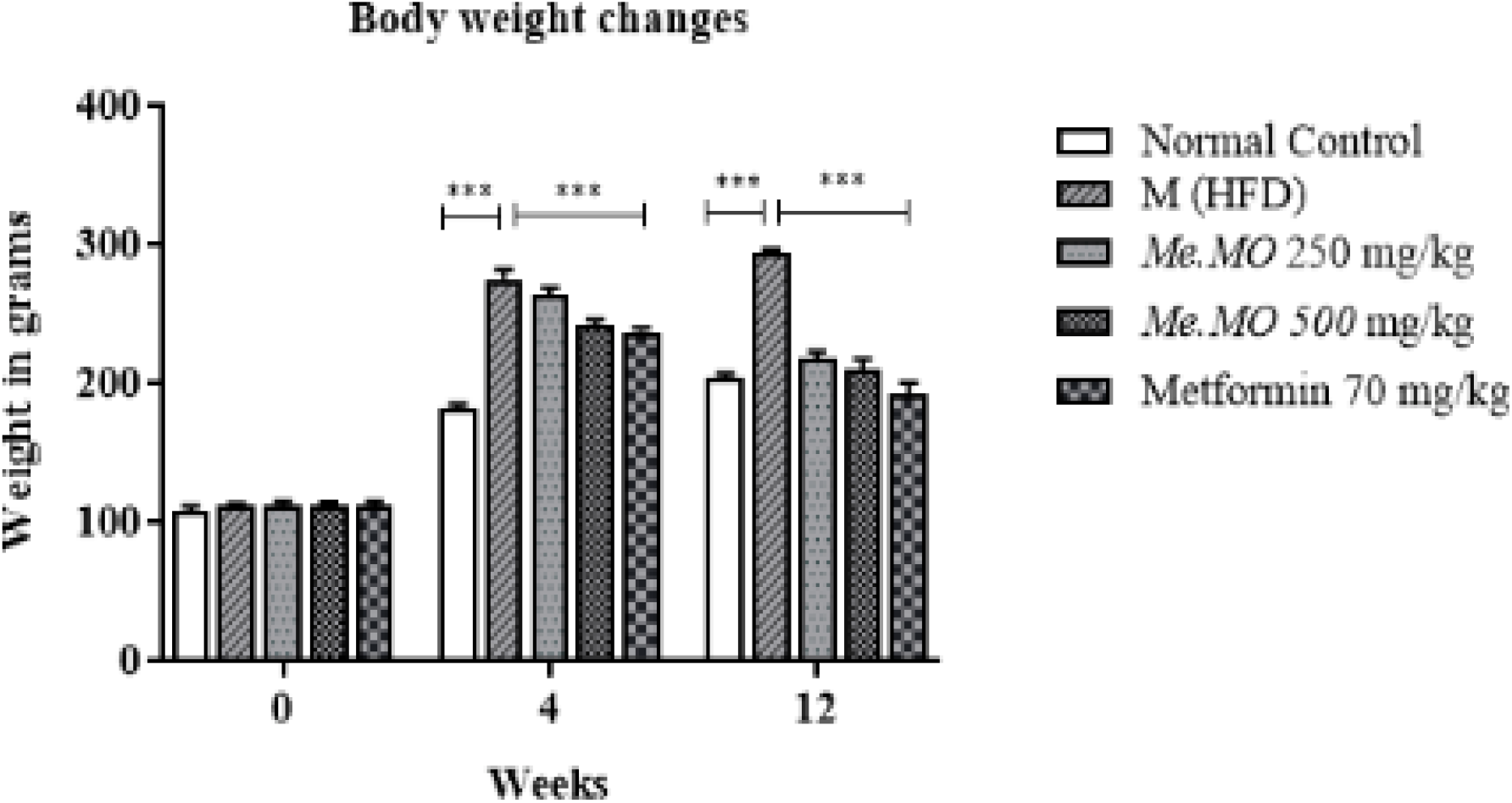
Shows the variation in body weights at 0, 4^th^ and 12^th^ week of study. The data shown represent the means ± SEM. \*\*\**P* < 0.01.

**Figure 4:**
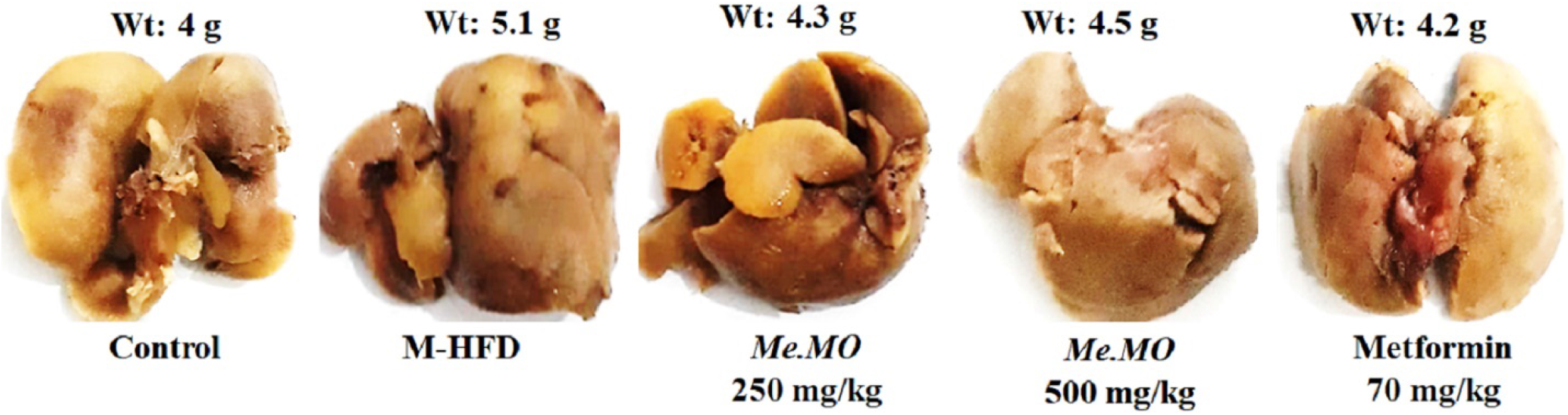
Macroscopic analysis and weight measurement of rats. The average liver weight of that group (n-=5) is given above the respective representative images.

### 3.3. Random blood glucose levels reversed by *Me*.*MO*

Data in figure 5A shows the effects of *Me*.*MO* on random blood glucose levels in animals. At zero week, the concentration of blood glucose was measured within the normal range with relative group comparability. Overall, the concentration of glucose was elevated in all groups due to the administration of HFD. On twelve-week, by comparing 250 mg/kg *Me*.*MO* (143.0 ± 0.40), 500 mg/kg *Me*.*MO* (157.0 ± 1.65) and metformin (111.0 ± 0.60) treatment groups with model-HFD (305.0 ± 0.50), significant (*p* < 0.001) decrease in blood glucose concentration was measured. However, control group animals (no treatment) showed normal glucose concertation throughout the study.

**Figure 5:**
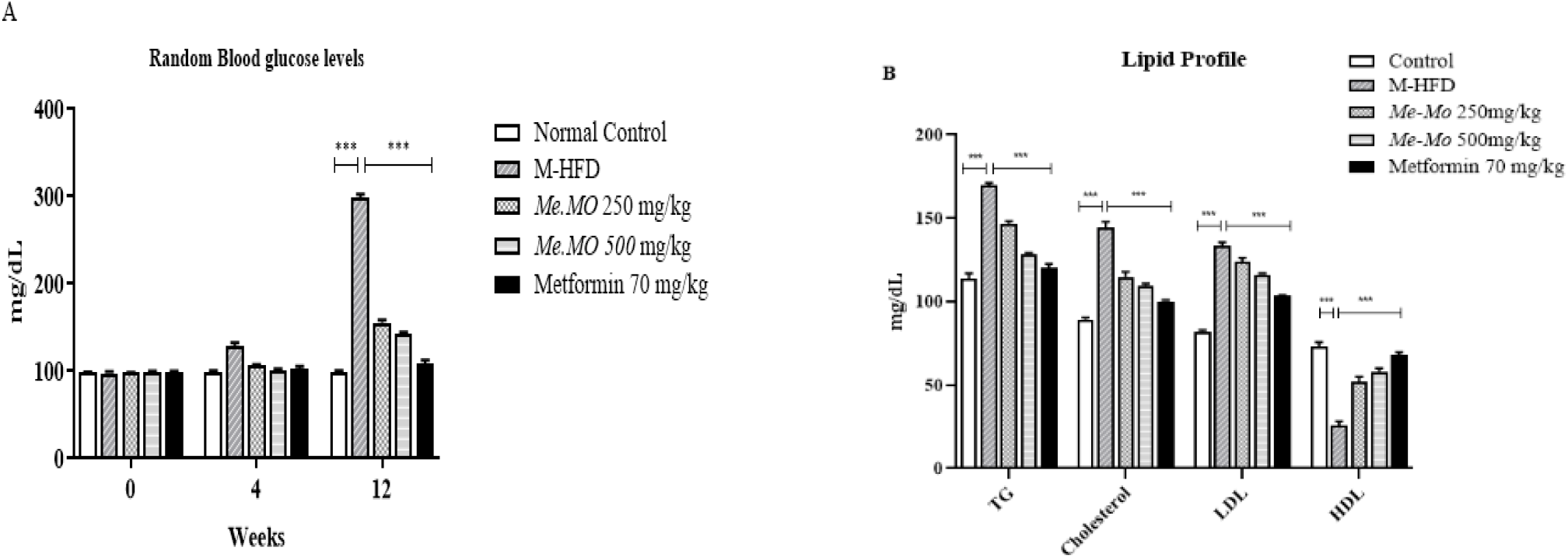
Section A shows graphical representation of random blood glucose levels in HFD and STZ induced model at 0, 4^th^ and 12^th^ week of treatment respectively. While section B shows graphical representation of lipid profile after twelve-week study. The data shown represent the means ± SEM. \*\*\**P* < 0.01.

### 3.4. Assessment of lipid profile

Data in figure 5B demonstrates a significant (*p* < 0.001) increment in cholesterol (145 ± 2.06), triglycerides (171 ± 4.35) and LDL (137±4.09) along with a significant decrease in HDL (36.3±2.90) levels in HFD induced rats. On the other hand, *Me*.*MO* treatment groups (250 and 500 mg/kg) showed significant (*p* < 0.001) reduction in serum cholesterol (112 ± 1.32, 112 ± 1.19), triglycerides (138 ± 3.14 and 143 ± 3.23) and LDL (114 ± 2.39 and 118 ± 3.20) concentrations respectively. However, *Me*.*Mo* raised the plasma HDL levels significantly. This elevation was significant (*p* < 0.001) in *Me*.*MO* treated groups as compared to model-HFD.

### 3.5. Histopathological analysis

Hepatic tissues showed fatty changes, hyperplasia, hypertrophy and macrophages inflammations in M-HFD and 500 mg/kg *Me*.*MO* treatment groups. While no specific changes were observed in 250 mg/kg *Me*.*MO* and metformin (70 mg/kg) treatment group except mild inflammation (green arrow). Moreover, relevant modifications in adipose tissues of model-HFD was also seen in form of adiponecrosis (blue arrow) (Figure 6).

**Figure 6:**
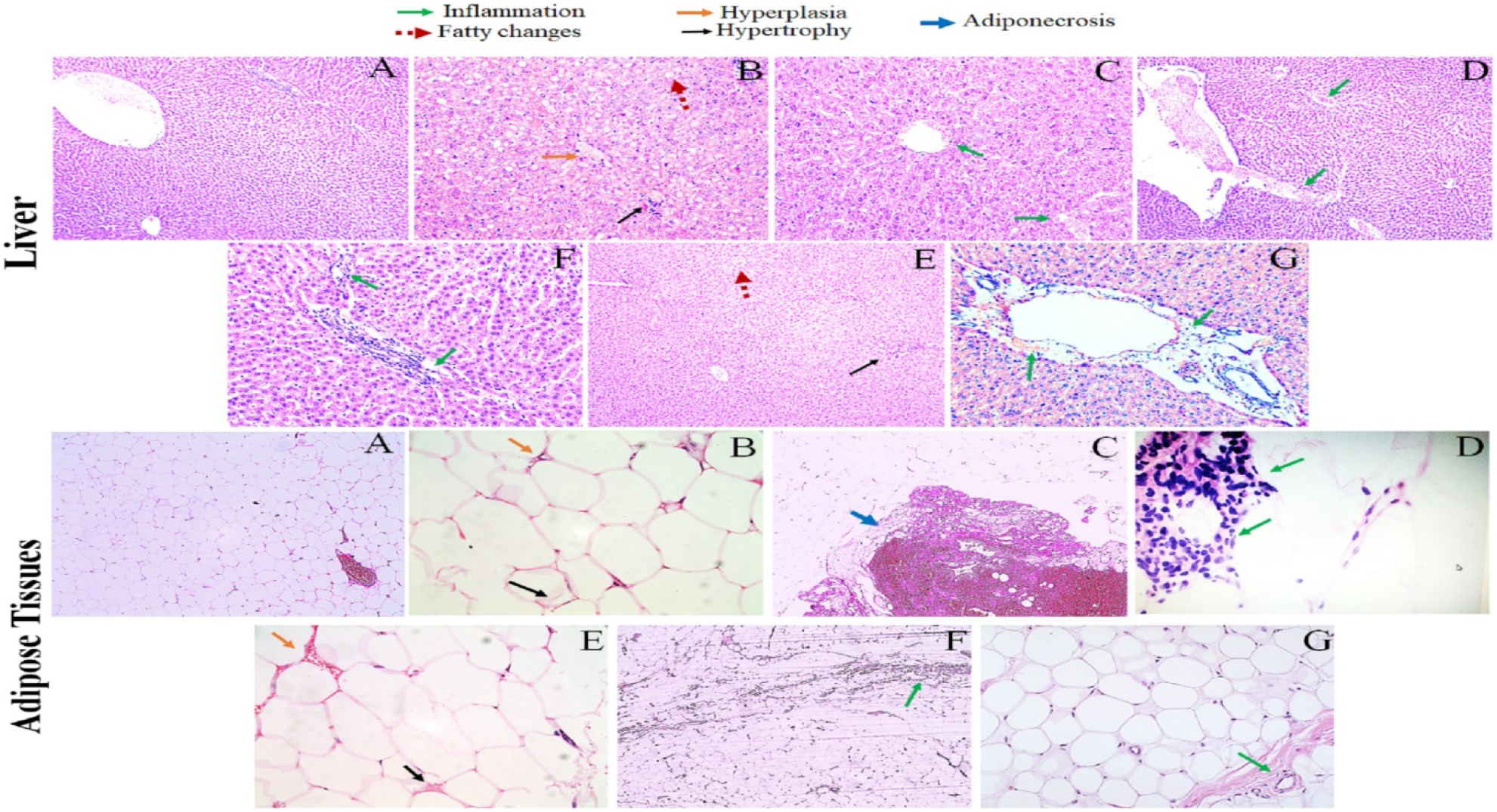
Histopathological analysis of HFD and STZ induced diabetic rats. A (Control), B and C (M-HFD), D (250 *Me*.*MO*), E and F (500 *Me*.*MO*) and G (Metformin 70 mg/kg).

### 3.6. Measurement of pro- and anti-inflammatory cytokines

Data in figure 7 is represented the effect of *Me*.*MO* and metformin on inflammatory cytokines. Findings showed a notably (*p* < 0.001) decrease in levels of pro-inflammatory cytokine IL-1 in *Me*.*MO* and standard treatment groups as compared to M-HFD group. Moreover, experimental data of *Me*.*MO* and metformin treatment groups showed significantly comparable (*p* < 0.001) reduction in IL-1β and TNF-α concentrations than M-HFD treatment group. Further, by comparing *Me*.*MO* and metformin-treated animals with M-HFD animals, no significant (*p* > 0.05) difference was analyzed statistically in levels of interleukin-10. Additionally, the concentration of anti-inflammatory cytokines such as interleukin-6 and COX-2 was observed to be significantly higher in *Me*.*MO* and standard treatment groups as compared to M-HFD group.

**Figure 7:**
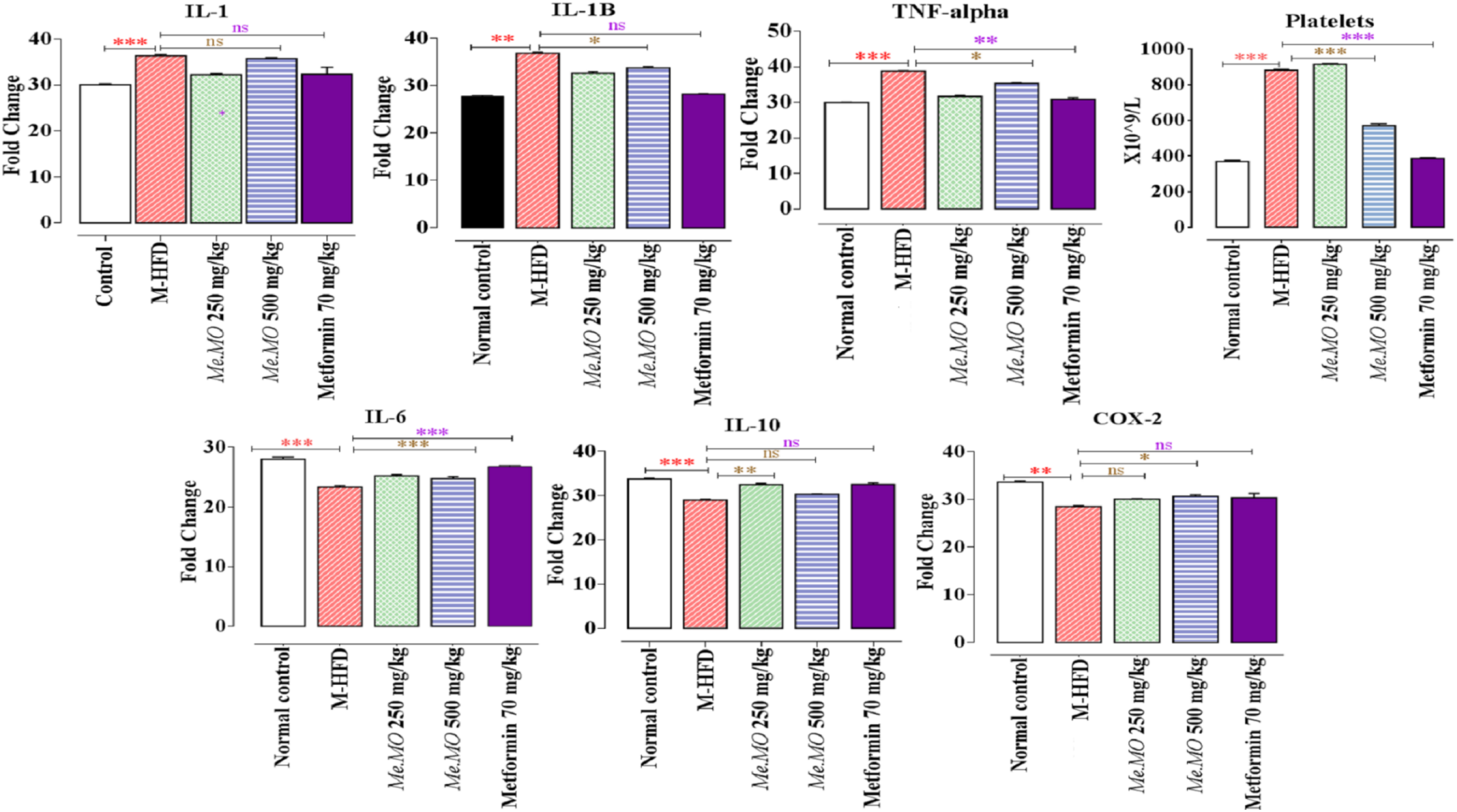
Graphical representation of random blood glucose levels in HFD and STZ induced diabetes mellitus model at 0, 4^th^ and 12^th^ week of treatment respectively. Red color stars show comparison of control with model-HFD while brown and violet color stars show comparison of model-HFD with *Me*.*MO* (250 and 500 mg/kg) and metformin treatment groups respectively. The data shown represent the means ± SEM. \*\*\**P* < 0.01.

## 4. Discussion

Weight gain and related fatty liver are considered early risk factors for much larger metabolic complications [28]. Obesity and accompanied fatty liver are regarded as benign states but often precede cardiovascular complications, chronic heart disease, stroke, type 2 diabetes, chronic inflammation and osteoarthritis [29]. The best-devised clinical strategy would be to counter the benign states (obesity and fatty liver) much earlier that avoid the major onslaught of destructive metabolic states. The current clinical regimen for these states are unable to provide the desired results or lead to adverse effects that make the therapy too unsuitable. Despite the widespread use of synthetic agents (such as antidiabetics, anti-hyperlipidemia and anti-obesity) treatment with herbal extracts has demonstrated more significant effects and safe drug profile in multiple pre- and clinical studies. Therefore, the development of innovative plant-based medicines that can mitigate obesity and accompanying complications are of higher priority for management [30].

The intake of HFD and STZ mimics the obese and high plasma glucose states with considerable inflammation [31, 32]. Obesity and related abnormalities can be prevented by consuming the plants extracts having an abundance of potential polyphenolic compounds [33]. In the present study, the result exhibited the provision of the promising therapeutic option of *Me*.*Mo* produced a remarkable decrease in body weight gain, fatty liver and related inflammation. Besides this, the accumulation of fats and insulin resistance in hepatic tissues could assume to be responsible for increasing rats liver weight in current research.

Moreover, lipid profiles including low-density lipoproteins (LDL), cholesterol and triglycerides are keynotes in the development of fatty deposits and plaques in vascular walls ultimately leading to a series of atherosclerosis, coronary heart disease and diabetes mellitus. As insulin has the inhibitory action on HMG-CoA reductase enzymes (primary contributor in cholesterol-rich LDL particles) thus, deficiency of insulin abnormally increases levels of free fatty acids and decreases their utilization in the body consequently developing diabetic conditions [34, 35]. On the other hand, high-density lipoproteins (HDL) play a primary contribution in lowering the risk of cardiovascular disease and stroke by scrubbing the inner walls of blood vessels from bad cholesterol [36]. The present work showed that in HFD-induced diabetic rats, *Me*.*MO* exerted inhibitory action on cholesterol, triglycerides, and LDL levels and cause upregulation of HDL concentration in the blood. These findings highlight the anti-obesity and lipid-lowering effects of *Me*.*MO* which are inconsistent with previous scientific studies [37].

Besides this, HFD induces significant insulin resistance and hyperinsulinemia in animals [38]. Similarly, in present research work oral administration of *Me*.*Mo* for consecutive 60 days significantly decreased HFD-induced glucose concentration, however, no marked changes were observed in the model-HFD group. The insulin-sensitizing effect of *Me*.*MO* may be due to stimulatory action on receptor insulin signaling pathways as attributed in previously published inline studies [39, 40].

Chronic inflammatory stress is the fundamental feature of obesity [41]. The inflammatory mediators serve as a magnet in attracting more fats to tissues that can store them. In turn, inflammatory cells themselves undergo metabolic rewiring in response to excess lipids [42] and often turn more active favoring autoimmune disease [43-46]. In the present work, results of RT-PCR analysis demonstrated that treatment with *Me*.*MO* for continuous two months decreased gene expression of chemo-cytokines. Therefore, it is speculated that *Me*.*MO* can prevent induction and propagation of inflammatory mediators release and have beneficial anti-inflammatory spectrum. In addition, histopathological analysis of liver and adipose tissues of *Me*.*MO* and metformin-treated diabetic rats showed a definitive reduction in adipocytes hypertrophy/hyperplasia and microvascular fatty changes with mild lymphocytes recruitment and no necrotic/apoptotic bodies. However, in M-HFD group adipocytes hyperplasia, fatty bodies, adiponecrosis and inflammatory infiltration was clearly seen.

HPLC analysis confirmed that *Me*.*MO* extract contains high content of polyphenolic compounds which have well-documented metabolic effects [47]. Quercetin plays a vital role to attenuate symptoms of the metabolic disease by decreasing oxidative stress and associated inflammation [48, 49]. Vanillic acid cut down hyperinsulinemia and improves antioxidants status in the body resultantly reducing the risk of cardiac problems [50]. Chlorogenic acid plays a dual role not only blocking the expression of glucose-6-phospatse but enhancing glucose uptake by skeletal muscles as well. Consequently, alleviates insulin sensitivity and glucose tolerance and dyslipidemia [51]. Synirgic acid downregulates altered levels of metabolic enzymes and stimulates β-cells regeneration to prevent pancreatic damage [52]. m-coumaric acid inhibits the overproduction of reactive oxygen species thus suppressing hyperglycemia-induced vascular damage [53]. Hence, in the present study, it can be attributed to the presence of these phytochemicals in *Me*.*MO* may be responsible for its anti-inflammatory and anti-obesity effects. We also believe that anti-inflammatory activities are responsible for lowering lipid load in experimental rats.

## 5. Conclusion

This study revealed that *Me*.*Mo* ameliorated body weights and related liver weight gains in HFD induced model. Interestingly, the reduction in body and liver weights was accompanied with inflammatory regulation. The study highlights the possible role of the immunometabolic axis in the manifestation of the anti-obesity potentials of *Me*.*Mo*.

## Authors’ contributions

Dr Nihal conceptualized the research topic. Dr Manal designed the study. All the authors provided valuable comments and suggestions on the study design. Dr El Shaima analyzed and interpreted the data. Dr Murtaza guided the statistical analysis. All the authors contributed in writing the manuscript and were responsible for revision. All authors read and approved the final manuscript.

## Acknowledgements

The study was supported by a research grant from the Deanship of Research & Graduate Studies, Ajman University # 2023-IRG-PH-11.

## Availability of data and materials

The datasets used and/or analyzed during the current study are available from the corresponding author upon reasonable request.

## Ethics approval and consent to participate

This study adhered strictly to the guidelines outlined in the Guide for Research Ethical Committee. The protocol received approval from the University’s Committee on the Ethics of Animal Experiments (reference number: A-H-F-8 Jun).

## Consent for publication

Not applicable.

